# Neighbor predation linked to natural competence fosters the transfer of large genomic regions in *Vibrio cholerae*

**DOI:** 10.1101/618009

**Authors:** Noémie Matthey, Sandrine Stutzmann, Candice Stoudmann, Nicolas Guex, Christian Iseli, Melanie Blokesch

## Abstract

Natural competence for transformation is a primary mode of horizontal gene transfer (HGT). Competent bacteria are able to absorb free DNA from their surroundings and exchange this DNA against pieces of their own genome when sufficiently homologous. And while it is known that transformation contributes to evolution and pathogen emergence in bacteria, there are still questions regarding the general prevalence of non-degraded DNA with sufficient coding capacity. In this context, we previously showed that the naturally competent bacterium *Vibrio cholerae* uses its type VI secretion system (T6SS) to actively acquire DNA from non-kin neighbors under chitin-colonizing conditions. We therefore sought to further explore the role of the T6SS in acquiring DNA, the condition of the DNA released through T6SS-mediated killing versus passive cell lysis, and the extent of the transfers that occur due to these conditions. To do this, we herein measured the frequency and the extent of genetic exchanges in bacterial co-cultures on competence-inducing chitin under various DNA-acquisition conditions. We show that competent *V. cholerae* strains acquire DNA fragments with an average and maximum length exceeding 50 kbp and 150 kbp, respectively, and that the T6SS is of prime importance for such HGT events. Collectively, our data support the notion that the environmental lifestyle of *V. cholerae* fosters HGT and that the coding capacity of the exchanged genetic material is sufficient to significantly accelerate bacterial evolution.

**Significance Statement:** DNA shuffled from one organism to another in an inheritable manner is a common feature of prokaryotes. It is a significant mechanism by which bacteria acquire new phenotypes, for example by first absorbing foreign DNA and then recombining it into their genome. In this study, we show the remarkable extent of the exchanged genetic material, frequently exceeding 150 genes in a seemingly single transfer event, in *Vibrio cholerae*. We also show that to best preserve its length and quality, bacteria mainly acquire this DNA by killing adjacent, healthy neighbors then immediately absorbing the released DNA before it can be degraded. These new insights into this prey-killing DNA acquisition process shed light on how bacterial species evolve in the wild.

## Introduction

The causative agent of the diarrheal disease cholera, *Vibrio cholerae*, is responsible for seven major pandemics since 1817, one of which is still ongoing. Due to its ability to rapidly spread in contaminated water, cholera poses a serious world health risk, affecting between 1 and 4 million people and causing 21,000–143,000 deaths per year, especially in poor or underdeveloped countries (1). Many disease-causing bacteria have developed mechanisms for rapidly evolving in response to environmental pressures, and these rapid changes are often responsible for the formation of new serogroups with pandemic potential. One way in which *V. cholerae* acquire new phenotypes is through horizontal gene transfer (HGT), which is the direct movement of DNA from one organism to another. A major mode of HGT is natural competence for transformation in which bacteria are able to absorb free DNA from their surroundings using their competence-induced DNA-uptake complex (2–4). When sufficient homology is present between the incoming DNA and the bacterial genome, the absorbed genetic material can be integrated into the genome via double homologous recombination at the expense of the initial DNA region. As an example of the significant power of this natural competence for gene uptake, we previously witnessed the gain of an ~40 kbp O139-antigen cluster at the expense of the original ~30 kbp O1-antigen cluster through natural transformation (followed by strong selective pressure exerted by antibiotics or phages; (5)), which significantly changed the phenotypes of these bacteria. And while Griffith’s experiment in 1928 unambiguously proved that transformation contributes to evolution and pathogen emergence, the general prevalence of non-degraded DNA with sufficient coding capacity has been questioned (6), drawing inquiries as to whether this mode of HGT could be responsible for the major changes causing pandemic strains to emerge.

The induction of competence in *V. cholerae* is tightly regulated (recently reviewed by (7)). Briefly, upon growth on the (molted) chitin-rich exoskeletons of zooplankton (8), the most abundant polysaccharide in the aquatic environment and therefore an important carbon source for chitinolytic bacteria (9), the expression pattern of *V. cholerae* is altered (10) to render it naturally competent for genetic transformation (11). Initially, when chitin degradation products are sensed by *V. cholerae*, it produces the regulatory protein TfoX (10–15). This competence activator positively regulates the expression of the major DNA-uptake machinery in the cell (4), providing a direct connection between growth on chitin and competence activation. Apart from TfoX, natural competence and transformation also depend on the master regulator of quorum sensing, HapR, in two ways: i) HapR acts as repressor of *dns*, which encodes an extracellular nuclease that inhibits transformation (16); and ii) HapR together with TfoX co-activates the transcription factor QstR, which further represses *dns* as well as activates several DNA-uptake genes (17, 18).

While the chitin-induced DNA-uptake complex of *V. cholerae* is able to absorb DNA from the surrounding (19–23), environmental DNA is often heavily degraded and therefore short in size (24, 25). In addition, free DNA is thought to originate from dead and therefore less fit bacteria, which renders the coding part of such genetic material non-favorable for naturally competent bacteria (26). In line with these arguments, we recently showed that *V. cholerae* does not solely rely on randomly released environmental DNA. Instead, it actively acquires “fresh” DNA from healthy, living bacteria through kin-discriminatory neighbor predation (27), which, conceptually, also occurs in other naturally competent bacteria (28). Neighbor predation in *V. cholerae* is accomplished by a contractile injection system known as the type VI secretion system (T6SS) that transports toxic effector proteins into prey (29–32). Intriguingly, the T6SS of pandemic *V. cholerae* is exquisitely co-regulated with its DNA-uptake machinery in a TfoX-, HapR-, and QstR-dependent manner when the bacterium grows on chitin (27, 33, 34), which increases the chances of the competent bacterium to take up freshly released DNA compared to free-floating “unfit” DNA. Notably, this coupling of competence and type VI secretion is also conserved in several non-cholera vibrios (35).

In the current study, we determined the extent of the absorbed and chromosomally-integrated prey-derived DNA. Previous studies had scored transformation events in other naturally competent Gram-negative bacteria such as *Haemophilus influenzae, Helicobacter pylori*, and *Neisseria meningitidis* (36–38). These former studies, however, relied on the supplementation of large quantities of purified DNA (with up to 50 donor genome equivalents per cell (37)) at the peak of the organism’s competence program (36). Such an approach, however, neither recapitulates the natural onset of competence nor discloses the fate of the DNA that is released from dying cells. Thus, to address these points and to mimic natural settings, we determined the frequency and extent of DNA exchanges under chitin-dependent co-culture conditions of two non-clonal *V. cholerae* strains. We show that the DNA transfer frequency is significantly enhanced in T6SS-positive compared to T6SS-negative strains and that large genomic regions are transferred from the killed prey to the competent acceptor bacterium.

## Results and Discussion

### The T6SS fosters horizontal co-transfer events encompassing two selective markers

To compare the absorption of T6SS-mediated prey-derived DNA as opposed to environmental DNA (released through, for example, random lysis), we first scored the transformability of T6SS-positive (wild-type [WT] predator) and T6SS-negative (acceptor) *V. cholerae* strains, which would allow us to directly measure the contribution of the T6SS on gene uptake. These two strains were co-cultured with non-kin prey (donor) bacteria that were all derived from the environmental isolate Sa5Y (27, 39–41) and contained two antibiotic resistance genes in their genomes: 1) An *aph* cassette (Kan^R^), which was integrated in the *vipA* gene on the small chromosome (chr 2); and 2) a *cat* cassette (Cm^R^), which was inserted at variable distances from the *aph* cassette on the same chromosome or, alternatively, on the large chromosome (chr 1). As shown in Figure 1, the WT predator strain efficiently absorbed and integrated the prey-released resistance cassettes (*aph* or *cat*), while the transformation efficiency for the T6SS-defective acceptor strain was significantly reduced (by 97.8% and 99.2% for *aph* and *cat*,respectively) (Fig. 1*A*). Moreover, comparable frequencies were observed for both selective markers, suggesting that their acquisition does not significantly affect the strains’ fitness under non-selective conditions. We tested whether these transfer events were indeed competence-mediated and not based on other modes of HGT using a strain with a competence-related DNA import deficiency in that it lacked the competence protein ComEA that reels external DNA into the periplasm (21). This *comEA*-minus strain was never transformed under these predator-prey co-culture conditions, confirming that the gene transfer did depend entirely on natural competence.

**Figure 1:**
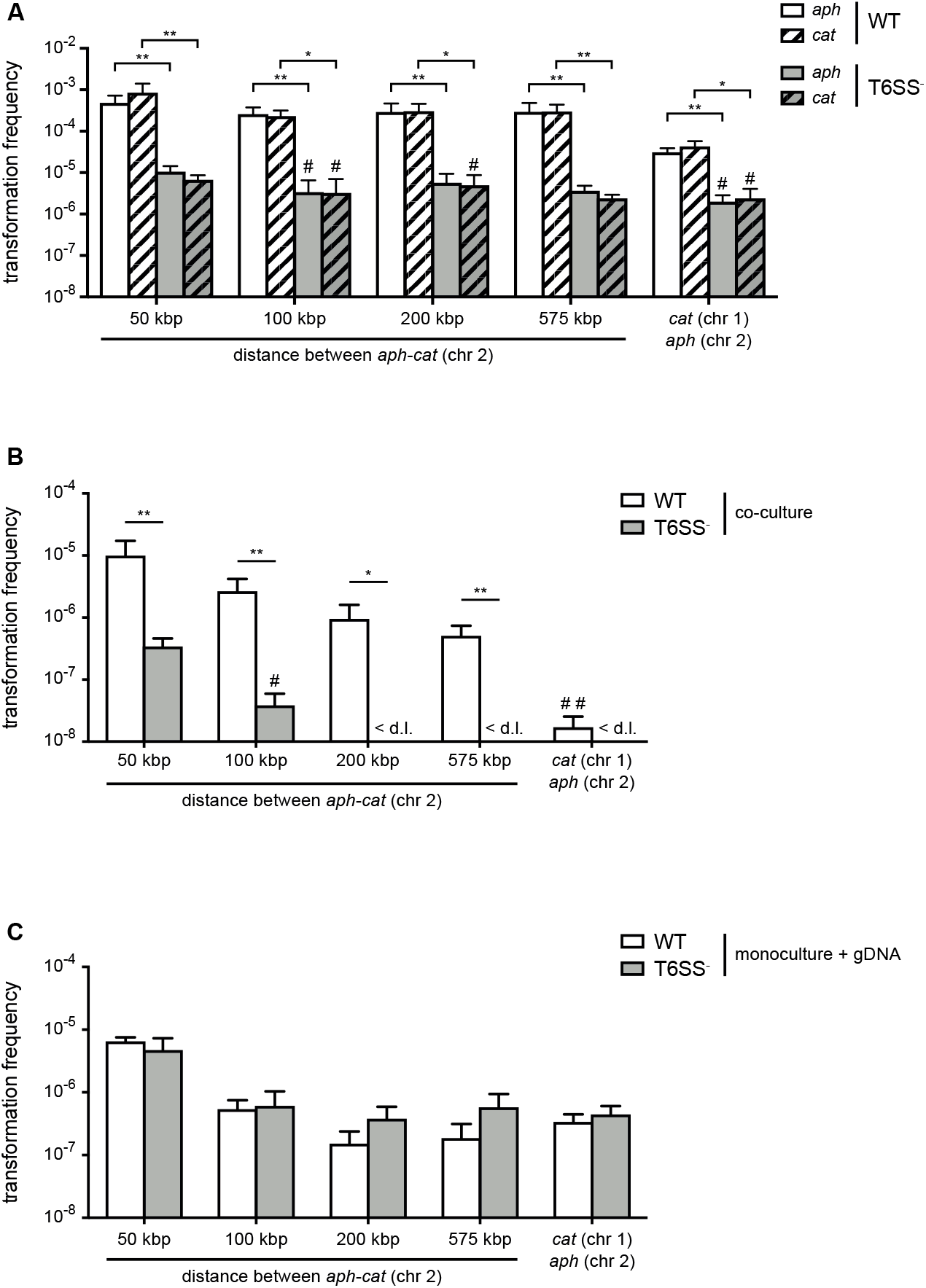
Type VI secretion system (T6SS) enhances horizontal gene transfer (HGT) of single- and double-resistance cassettes if carried *in cis*. (*A-B*) Transformation occurs in predator/prey co-cultures. To induce natural competence, the WT or a T6SS-negative derivative (A1552Δ*vasK*; T6SS^−^) was co-cultured on chitin with different prey strains (Sa5Y-derived) that carried two antibiotic resistance cassettes: *aph* in *vipA* (chr 2) and *cat* at variable distances from *aph* on the same chromosome or on chr 1, as indicated on the X-axis. Transformation frequencies (Y-axis) indicate the number of transformants that acquired (*A*) a single resistance cassette or (*B*) both resistance cassettes divided by the total number of predator colony forming units (CFUs). (*C*) Natural transformation is not impaired in the T6SS^−^ acceptor strain. Purified genomic DNA (gDNA) was added to competent WT or T6SS^−^ strains. (*A-C*) Data represent the average of three independent biological experiments (± SD, as depicted by the error bars). For values in which one (#) or two (##) experiments resulted in the absence of transformants, the detection limit was used to calculate the average. <d.l., below detection limit. Statistical significance is indicated (*p < 0.05; \*\**p* < 0.01).

Next, we scored the frequencies of transformants that had adopted resistance against both antibiotics, which would show the possibility of two transformation events or the transfer of a large piece of DNA (indicated by the distance between the two genes on the same chromosome). These transformations occurred, as expected, at lower rates compared to single-resistant clones and were mostly below the limit of detection for the T6SS-minus acceptor strain (Fig. 1*B*).Interestingly, we observed a gradual decrease in the frequencies the further the two resistance genes were apart from each other on the same chromosome, while a sharp drop occurred in the number of recovered transformants when the two resistance genes were carried on the two separate prey chromosomes (Fig. 1*B*). While the latter scenario unambiguously requires at least two separate DNA-uptake events, the former, in which the resistance markers are carried *in cis*,could reflect a mix between single and multiple DNA absorption and integration events. When purified genomic DNA was instead provided as the transforming material to simplify the experiment and provide measurable results for all conditions, the *in cis* double-resistance acquisition efficiencies reached a comparable range to the *in trans* efficiencies when the two resistance genes were separated by at least 100 kbp. This suggested that the more efficient transformations of less than 100 kbp likely often occurred through a single acquisition (Fig. 1*C*). Furthermore, the WT predator and T6SS-minus acceptor behaved similarly when purified DNA was provided, which makes sense as the need for active DNA release through neighbor predation was eliminated. Based on these data and the fact that the double-acquisition rates for the T6SS-minus acceptor strain were mostly below the detection limit in the prey scenario, we hypothesized that neighbor predation might foster the transfer of long DNA stretches, which frequently exceeded 50 kbp and therefore carry significant coding capacity.

### Comparative genomics of pandemic strain A1552 and environmental isolate Sa5Y

To test our hypothesis that the T6SS contributes to the horizontal transfer of large DNA fragments, we used a whole-genome sequencing (WGS) approach to properly outline the transferred DNA regions. To do this using WGS, we first needed to characterize the genomes of both the predator/acceptor (A1552) and the prey/donor (Sa5Y) strains for which long-read PacBio sequencing data and *de novo* assemblies without further analysis were recently announced (41). A1552 is a pandemic O1 El Tor strain (42) belonging to the LAT-1 sublineage of the West-African South American (WASA) lineage of seventh pandemic *V. cholerae* strains (43) (see *SI Appendix* for details) while strain Sa5Y was isolated from the Californian Coast (39, 40). To understand their genomic arrangements, we also compared these strains to the reference sequence of *V. cholerae* (O1 El Tor strain N16961; (44)) and a re-sequenced laboratory stock of the latter. Details on the comparative genomics between the three pandemic strains (N16961 (44), the newly sequenced and *de novo*-assembled genome sequence of the laboratory stock of N16961, and A1552) are provided in the *SI Appendix* and as Figure S1-S2. We expected to see significant differences in the pandemic A1552 strain compared to the environmental isolate Sa5Y in terms of the absence/presence of genomic features and single nucleotide polymorphisms (SNPs) in core genes that would allow us to measure HGT events occurring between the strains, and several of these major differences are highlighted here. Indeed, as expected from its non-clinical origin, the environmental isolate lacked several genomic regions, including those that encode major virulence features, namely *Vibrio* pathogenicity islands 1 and 2 (VPI-1, VPI-2), *Vibrio* seventh pandemic islands I and II (VSP-I, VSP-II (45)), the cholera toxin prophage CTX (46), and the WASA-1 element. In addition, the strain’s O-antigen cluster differed significantly from the O1-encoding genes of pandemic strain A1552 (Fig. 2). The region that differed the most between both strains was the integron island, which is consistent with the role of this assembly platform in fostering the incorporation of exogenous open reading frames (47). Given these major differences between strain A1552 and Sa5Y and, in addition, an overall SNP frequency of approximately 1 in 55 nucleotides for conserved genes, we concluded that HGT events occurring between these two strains on chitinous surfaces could be precisely scored using short-read sequencing. Apart from this important genomic information, we also noted that the pandemic strains as well as Sa5Y contained previously unrecognized rRNA operons, with nine or ten rRNA clusters in total compared to the initially reported eight (44).

**Figure 2:**
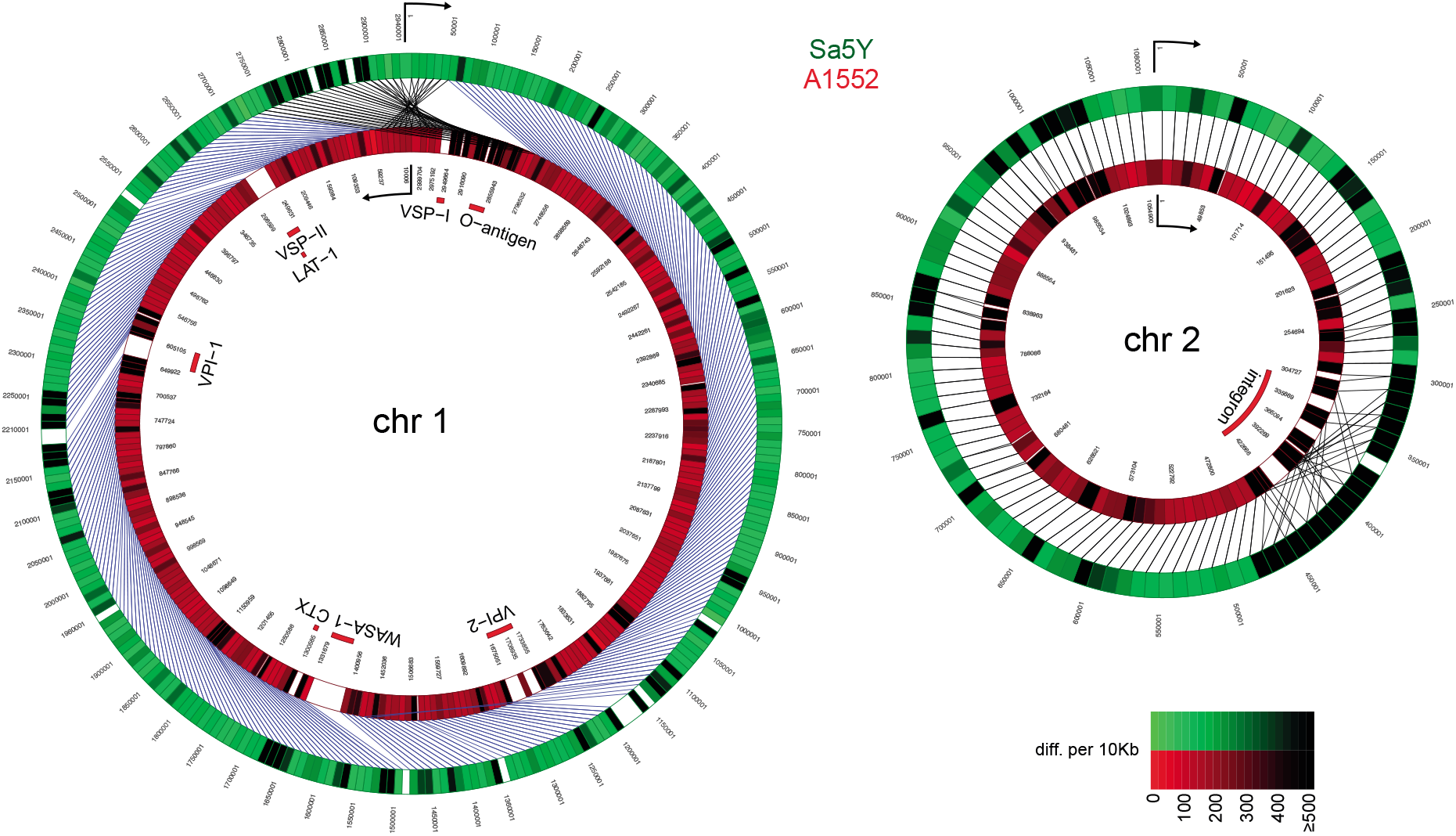
Comparative genomics of pandemic strain A1552 and the environmental isolate Sa5Y. The genomic sequences of chr 1 and 2 of Sa5Y (green) were segmented in 10-kbp-long fragments and aligned against the respective chromosome of the reference A1552 (red). To simplify visualization, chr 1 of strain A1552 was inverted and plotted counter-clockwise relative to Sa5Y (due to the large inversion in this strain; *SI Appendix*), as indicated by the arrow. To represent the differences between the two genomes, a color intensity scale was used that corresponded to the number of differences (SNP or indel), from 0 to ≥500 as measured per 10 kbp fragment. White regions show no homology. Important genomic features of pandemic *V. cholerae* are highlighted inside the rings.

### Released DNA from T6SS-killed prey leads to the transfer of large genomic regions

As our previous study witnessed gene transfers between *V. cholerae* bacteria (5) though neither scored the full extent of the transferred DNA region nor took T6SS-mediated neighbor predation into consideration, we sought to next determine how much genetic material would be absorbed and integrated by competent *V. cholerae* upon neighbor predation. To do this, we co-cultured the predator (A1552) and prey (Sa5Y) strains on chitinous surfaces for 30 h without any deliberate selection pressure. To be able to afterwards screen for the transfer of at least one gene, we first integrated an *aph* cassette within the *vipA* of strain Sa5Y, which concomitantly deactivated the prey’s T6SS, to select kanamycin-resistant transformants of strain A1552. Using this system, resistant transformants of A1552 were selected at an average frequency of 1.8 x 10^−4^ after the 30 h co-culturing on chitin (Fig. S3), and 20 of those transformants were randomly picked for further analysis. After three independent experiments, the whole genome of each of the 60 transformants was sequenced, and the reads were mapped to either the predator’s or the prey’s genome sequence (see *SI Appendix* for detailed bioinformatic analysis). As shown in Figure 3, apart from the common acquisition of the *aph* resistance cassette, the location and the size of the prey-donated genomic region differed significantly between most transformants. Previous estimates of the average length of total acquired DNA were made in experiments using purified donor gDNA and were considered to be ~23 kbp (40). Importantly, we observed in these new experiments that the average length of the total acquired DNA, meaning the DNA surrounding the *aph* cassette plus any transferred regions elsewhere on either of the two chromosomes (Fig. S4), was almost 70 kbp and therefore significantly larger than the previous estimates. Around 15% of all transformants acquired and integrated more than 100 kbp (Fig. 3*B*), which was previously considered unlikely due to absence of such long DNA fragments in the environment. Consistent with the principle of natural transformation, it should be noted that the new DNA was acquired through double homologous recombination such that it replaced the initial DNA region and the overall genome size did not significantly change. Further analysis indicated that about 50% of the strains experienced a single HGT event around the *aph* cassette, while the others exchanged regions in up to eight different locations on the two chromosomes (Fig. 3*C*). Finally, we analyzed the length of continuous DNA stretches that were acquired from the prey and observed that those ranged from a few kbp up to 168 kbp (Fig. 3*D*). Collectively, these data indicate that *V. cholerae* can acquire large genomic regions from killed neighbors with an average exchange of more than 50 kbp or ~50 genes. This finding contradicts the notion that natural transformation cannot serve for DNA repair or acquisition of new genetic information due to the insufficient length and coding capacity of the acquired genetic material.

**Figure 3:**
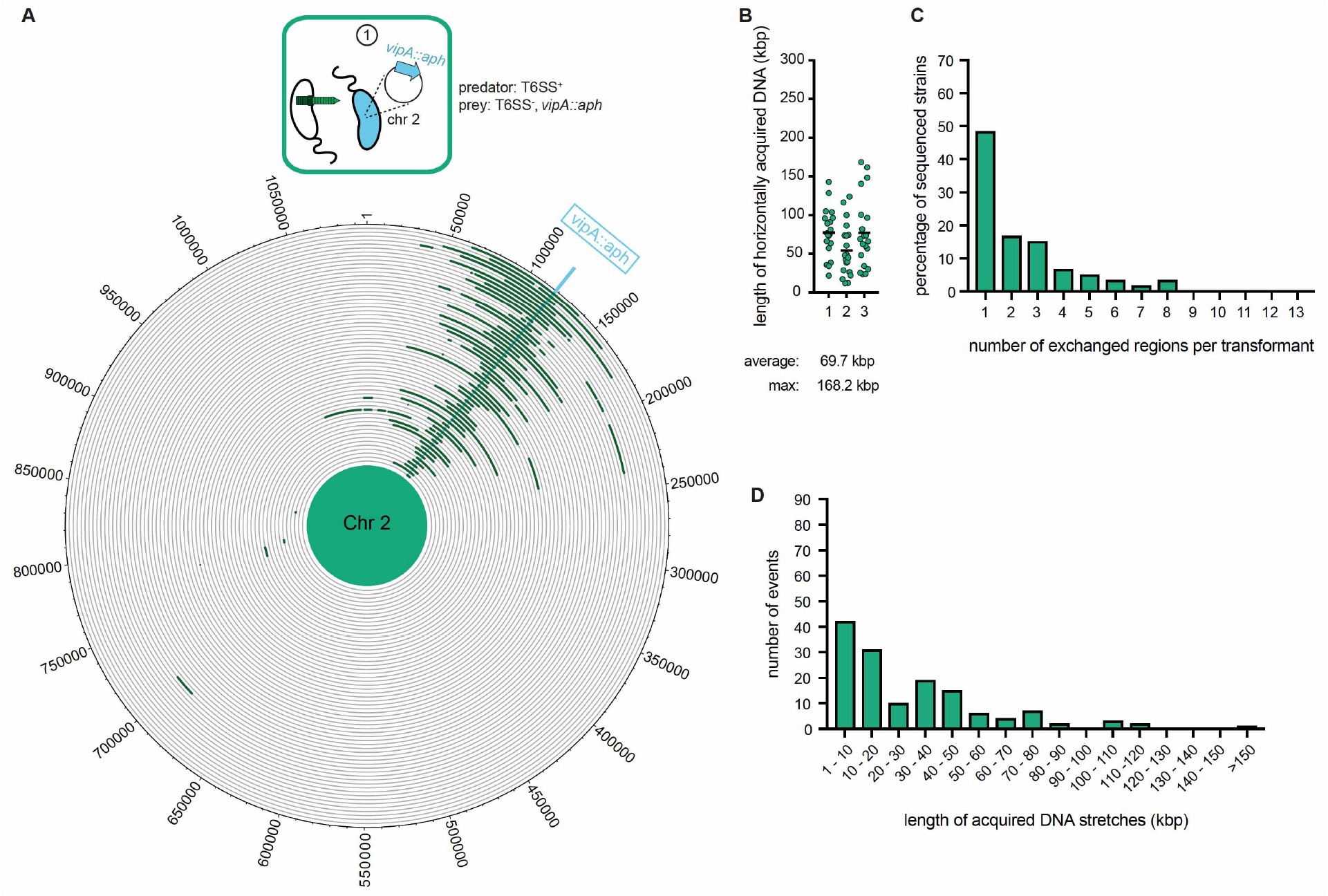
Whole-genome sequencing (WGS)-based quantification of horizontally acquired DNA. WGS analysis of transformants after prey killing and DNA transfer. Twenty kanamycin-resistant transformants were selected per independent biological experiment (n = 3). (*A*) The scheme represents the experimental setup of the co-culture experiment (condition ①). Sequencing reads for each transformant were mapped onto the prey genomes to visualize the transferred DNA regions (in dark green; see Fig. S4 for both chromosomes). The position of the resistance cassette (*aph*) is indicated by the light blue line. (*B*) Total DNA acquisition frequently exceeds 100 kb. The total length of horizontally acquired DNA is indicated on the Y-axis for each transformant. Data are from three biologically independent experiments as indicated on the X-axis. Average and maximum lengths are indicated below the graph. (*C*) Multiple transferred DNA regions were identified in the transformants. Percentage of transformants (n = 60) that exchanged one or more DNA regions, as indicated on the X-axis. (*D*) Large DNA stretches are transferrable by transformation. The length of individual consecutive DNA stretches was determined as indicated on the X-axis.

### Transformation by purified DNA only occurs if correctly timed

To better understand the DNA acquisition and integration potential of naturally competent *V. cholerae*, we next compared the data described above, which we refer to from now on as condition ① using experiments varying the aspects of neighbor predation and DNA supplementation (Fig. 4*A*). First, the acceptor strain was grown in a monoculture immediately supplemented with purified genomic DNA (gDNA) derived from the same donor (prey) strain as described above. Notably, when the gDNA was added at the start of the chitin-dependent culture, no transformants were reproducibly detected from three independent biological experiments, suggesting that free DNA is rapidly degraded under such conditions. This finding is consistent with our previous work in which we demonstrated that *V. cholerae* produces an extracellular and periplasmic nuclease Dns (16, 20) that degrades transforming material. At high cell density (HCD), where competence is induced, *dns* is partially repressed through direct binding of HapR (16, 17), and this repression is reinforced by the transcription factor QstR (17, 18). We therefore concluded that the simultaneous expression of both machineries, concomitantly with a strong repression of *dns*, is a prerequisite for successful DNA transfer. Indeed, such coordinated expression would ensure that T6SS-mediated attacks are exquisitely timed with low nuclease activity so that the prey-released DNA can be efficiently absorbed.

**Figure 4:**
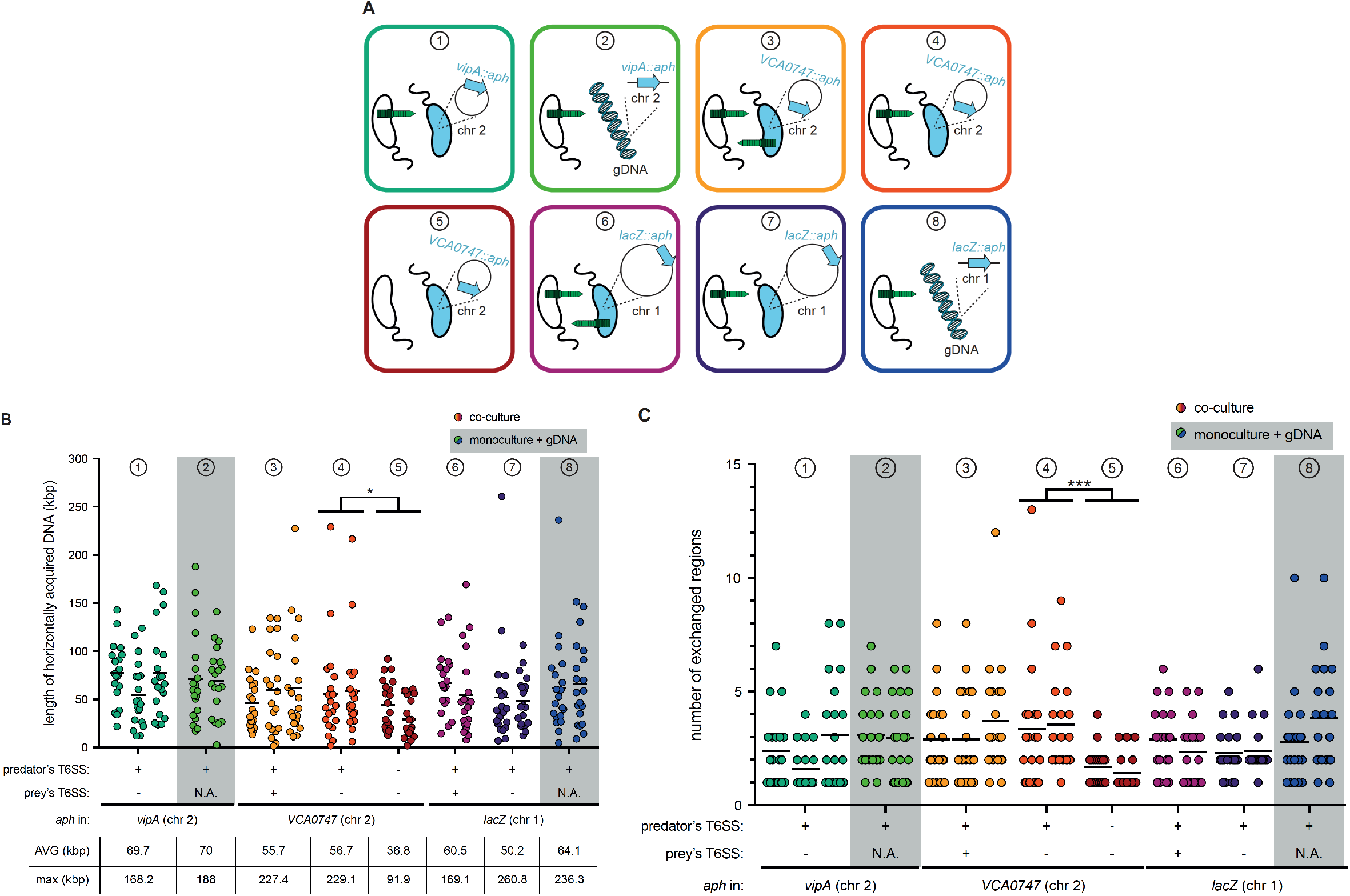
T6SS-mediated neighbor predation followed by DNA uptake enhances the frequency and length of transferred DNA stretches. (*A*) Scheme representing the eight experimental conditions tested in this study. Each scheme indicates whether the transformants acquired the *aph* resistance gene from a prey bacterium (blue) (position of *aph* indicated on the zoomed-in circles of chr 1 or chr 2) or from purified genomic DNA (gDNA). In the former case, the killing capacity of the predator (white) and prey (blue) is shown by the presence or absence of the dark green T6SS structure. The same color code is maintained throughout all figures. (*B-C*) Transformants from independent biological experiments (n ≥ 2) were analyzed by WGS for each of the conditions ①-⑧, as indicated at the top of each graph. The main features of predator and prey/gDNA are summarized below the X-axis. Panels (*B*) and (*C*) depict the total length of acquired DNA and the number of exchanged DNA stretches, respectively, for each transformant. N.A., not applicable. Statistical analysis is based on a pairwise comparison between different conditions. * *p* < 0.05, *** *p* < 0.001.

As we previously showed that the addition of purified gDNA after ~20–24 h of growth on chitin wasn’t prone to degradation by Dns (48), we next choose this time point to probe the DNA acquisition capability using purified DNA (condition ②; Fig. 4*A*). Doing so led to similar transformation frequencies as those observed for the prey-released DNA caused by T6SS attacks (condition ①; Fig. S3*A*). WGS of 20 transformants from two biologically independent experiments likewise resulted in similar DNA acquisition patterns with average and maximum DNA acquisitions of 70 kbp and 188 kbp, respectively, and the presence of multiple exchanged regions of varying sizes (Fig. 4 and Fig. S5). While we cannot entirely exclude that the maximum length of individual DNA stretches was biased by the purification step, despite the fact that we chose a method that was designed for chromosomal DNA isolation of 20–150 kbp sized fragments (see methods), our results suggest that the maximum DNA acquisition length of single fragments is probably reached between 100–110 kbp (Fig. S5). Moreover, the comparable acquisition patterns between conditions ① and ② (Fig. 4) imply that the prey-released DNA in condition ① is neither heavily fragmented nor is its accessibility or absorption by the competent acceptor bacterium significantly hindered due to, for example, DNA-binding proteins.

### Prey-exerted T6SS counter attacks do not change the DNA transfer pattern

Since the *aph* cassette was located within the T6SS sheath protein gene *vipA* in the above experiments, we wondered if this T6SS inactivation biased the DNA transfer efficiency. We therefore repeated the above-described experiments using prey strains that carried the *aph* cassette on the opposite site of chr 2 (within gene VCA0747; condition ③). As shown in Figure S3, similar transformation frequencies were observed independent of the position of the *aph* cassette. Moreover, WGS of 2 x 20 transformants showed similar average and maximum DNA acquisition values (55.7 kbp and 227.4 kbp, respectively; Fig. 4) as well as similar distribution patterns around the resistance marker (Fig. S6). However, while not statistically supported, it appeared as if these conditions were prone to the acquisition of multiple non-connected regions, as transformants with only single/connected exchanges dropped from ~50% (Fig. S4 for condition ①) to around 20% (Fig. S6 for condition ③). Based on this observation, we hypothesized that the now-restored T6SS-mediated killing capacity of the prey led to the additional release of genomic DNA from the predator, which interfered with the uptake of prey-released DNA. To test this idea, we repeated condition ③ (e.g., *aph* within VCA0747) though again inactivated the T6SS of the prey using a non-selected marker (*cat*; condition ④), expecting the results to be similar to those of condition ① if this hypothesis was correct. No statistically significant differences were observed between both conditions (③ and ④) for all tested characteristics including transformation frequency (Fig. S3B), number of exchanges, and separate and collective length (Fig. 4 and Figs. S6-S7), suggesting that predator-released DNA does not interfere with the predator’s overall transformability by the prey-released DNA. However, we acknowledge that the technical limitations of the experimental setup did not allow the identification of complete revertants that first acquired and then again lost the *aph* cassette.

### T6SS-independent prey lysis rarely triggers DNA transfer and results in shorter DNA exchanges

We next tested whether T6SS-mediated DNA release impacted the length of the exchanged region, which would support the above speculation that the intimate co-regulation of type VI secretion, nuclease repression, and DNA uptake ensures that freshly released DNA is rapidly absorbed by the predator and is therefore less prone to fragmentation. Such co-regulation would not hold true for T6SS-independent DNA release as a result of random cell lysis, so we tested the transfer efficiency of the *aph* cassette under conditions in which both donor and acceptor strains were T6SS-defective (condition ⑤; Fig. 4 and S8). Under such conditions, the transformation frequency dropped by 99.7% (Fig. S3*B*), and WGS of 2 x 20 of these rare transformants showed significant differences. Indeed, the average and maximal length of acquired DNA (Fig. 4*A*) and the number of exchanged regions (Fig. 4*B*) were significantly different when T6SS+ versus T6SS− acceptor strains were compared, with the latter exchanges never exceeding four events compared to up to 13 events for T6SS-mediated DNA release (Figs. S7 and S8). Based on these data, we conclude that T6SS-mediated DNA acquisition not only increases the transfer efficiency by ~100-fold but also fosters the exchange of multiple DNA stretches of extended lengths.

### T6SS-mediated DNA exchanges are not limited to the small chromosome

The experiments described above were designed to primarily score the transfer efficiency of DNA fragments localized on chr 2. The rationale behind this approach was a recent population genomic study on *Vibrio cyclitrophicus* that suggested the mobilization of the entire chr 2 and caused the authors to speculate: “how often and by what mechanism are entire chromosomes mobilized?” (49). In the current study, we were unable to experimentally show such large transfer events. We considered four potential reasons for the absence of such large transfers: 1) mild fragmentation of prey-released DNA that excluded fragments above ~200 kb; 2) limited DNA uptake and periplasmic storage capacity of the acceptor strain (20, 21); 3) limited protection of the incoming single-stranded DNA by dedicated proteins (such as Ssb and DprA; (50, 51)); or 4) lethality of larger exchanges due to the presence of multiple toxin/antitoxin modules within the integron island on chr 2 of *V. cholerae* (52). While technical limitations did not allow us to address the first three points, we followed up on the last idea by repeating the above-described experiments using prey strains in which the *aph* cassette was integrated on the large chromosome 1 (inside *lacZ*). We used these to test three (co-)culture conditions in which the prey strain was either T6SS-positive (condition ⑥), T6SS-negative (condition ⑦), or replaced by purified gDNA (condition ⑧; Fig. 4A). As shown in Figure S3, the *aph* cassette was again transferred with high efficiency from the killed prey strain to the acceptor strain. However, comparing conditions ⑥ (co-culture conditions) and ⑧ (prey-derived purified gDNA as transforming material) revealed a small but significant transformation increase (~ 4-fold; Fig. S3*A*). Based on these data, we speculate that the larger size of chr 1 (~3 Mb) compared to chr 2 (~1 Mb) slightly lowers the probability of acquiring the *aph* cassette when released from killed prey. This effect becomes negligible when purified gDNA is provided, most likely due to the size constraints of the purification procedure (max. 150 kb). Consistent with this idea was the finding that purified gDNA from all those prey strains described in this study resulted in the same level of transformation no matter where the selective marker was located (Fig. S3*D*).

Next, we randomly picked 20 transformants from two biologically independent experiments for each of these three experimental conditions (⑥ to ⑧) and sequenced their genomes (Fig. S9-S11). The analysis of these transformants showed that the average and maximum DNA acquisition values were highly comparable to those described above for DNA exchanges on chr 2 (Fig. 4*A*) and that multiple exchanged regions were likewise observed (Fig. 4*B*). We therefore conclude that prey-derived transforming DNA can equally modify both chromosomes. Moreover, our data suggest that consecutive stretches of exchanged DNA above ~200 kbp either do not occur or occur at levels below the detection limit of this study, and that this size limitation is not caused by the toxin/antitoxin–module-containing integron island on chr 2.

## Conclusion

Based on the data presented above, we conclude that T6SS-mediated predation followed by DNA uptake leads to the exchange of large DNA regions that can bring about bacterial evolution. This finding is consistent with the heterogeneous environmental *V. cholerae* populations that were observed in cholera-endemic areas (53). Still, an open question that remains is why pandemic cholera isolates are seemingly clonal in nature (43, 54–56), and we propose two explanations for this. First, sampling strategies might be biased for the selection of the most pathogenic strains and, concomitantly, exclude less virulent variants that have undergone HGT events. Secondly, transformation-inhibiting nucleases similar to Dns (16) have recently spread throughout pandemic *V. cholerae* isolates as part of mobile genetic elements (experimentally shown for VchInd5 (57) and predicted for SXT (58)), which makes these pandemic strains less likely to undergo HGT events. One could also argue that pandemic *V. cholerae* are rarely exposed to competence-inducing chitinous surfaces due to the prevalence of inter-household transmission throughout cholera outbreaks (59). Yet *in vivo*-induced antigen technology (IVIAT) assays showed strong human immune responses against proteins of the DNA-uptake pilus that fosters natural transformation, kin recognition, and chitin colonization (10, 11, 19, 23), which contradicts this idea. Indeed, the major pilin PilA was most frequently identified by IVIAT together with the outer-membrane secretin PilQ (60), which suggests that the bacteria encounter competence-inducing conditions either before entering the human host or after its colonization. The latter option is not, however, supported by *in vivo* expression data from human volunteers (61). Notably, our work shows the incredible DNA exchange potential that chitin-induced *V. cholerae* strains exert under co-culture conditions and future studies are therefore required to better understand strain diversity in clinical and environmental settings in the absence of sampling biases.

## Materials and Methods

### Bacterial strains, plasmids, and growth conditions

The bacterial strains and plasmids used in this study are described in *SI Appendix*, Table S1. Unless otherwise stated, bacteria were grown aerobically in LB medium under shaking conditions or on solid LB agar plates (1.5% agar). Growth on chitinous surfaces was performed as previously described (27, 48). Additional details are provided in the *SI Appendix*.

### Preparation of genomic DNA

Genomic DNA (gDNA) was purified from a 2 ml culture of the respective strain. DNA extraction was performed using 100/G Genomic-tips together with a Genomic DNA buffer set as described in the manufacturer’s instructions (Qiagen). After precipitation, the DNA samples were transferred into Tris buffer (10 mM Tris-HCl, pH 8.0). This was preferred over rapid gDNA isolation kits such as the DNeasy Blood & Tissue kit (Qiagen), as the latter isolation kit is strongly biased towards shorter DNA fragments (predominantly 30kb in length compared to up to 150kb for the 100/G columns, as stated by the manufacturer).

### Natural transformation assay

Natural transformation assays were performed by adding purified gDNA to the chitin-grown bacteria or by co-culturing the two non-clonal *V. cholerae* strains. To set up the experiments, the bacterial strains were grown as an overnight culture in LB medium at 30°C. After back dilution, the cells were incubated in the presence of chitin flakes (~80 mg; Sigma-Aldrich) submerged in half-concentrated (0.5x) defined artificial seawater medium (11). When purified DNA served as the transforming material, 2 μg of the indicated gDNA was added after 24 h of growth on chitin, and the cells were incubated for another 6 hours. At that point, the bacteria were detached from the chitin surfaces by vigorous vortexing and then were serially diluted. Colony-forming units (CFUs) were enumerated on selective (antibiotic-containing) or non-selective (plain LB) agar plates, and the transformation frequency was calculated by dividing the number of transformants by the total number of CFUs. For mixed community assays, the two strains were inoculated simultaneously at a ratio of 1:1. These mixtures were incubated for 30 h before the bacteria were harvested, diluted, and plated, as described above. All transformation frequency values are averages of three biologically independent experiments except for WGS conditions ② and ④–⑧, wherein the averages of two independent experiments are depicted.

### Whole-genome sequencing

For WGS, transformation assays were performed as described above using eight different experimental conditions (Fig. 4 and listed in *SI Appendix*, Table S2). To focus on the acquisition potential of strain A1552, conditions ⑤-⑧ used transformation-deficient prey strains (e.g., Sa5Y derivatives in which a *bla* cassette interrupted the DNA translocation channel protein encoding gene *comEC* (19)). The 360 recovered transformants (3 x 20 for experimental conditions ① and ③, which showed high levels of reproducibly, followed by 2 x 20 for all other conditions; see *SI Appendix*, Table S2 for details) were grown overnight in LB medium. Genomic DNA extraction was performed as described above. Further processing of the samples was conducted by Microsynth (Balgach, Switzerland). The quality of the DNA samples was verified before DNA libraries were prepared using a Nextera XT Library Prep kit (Illumina). Paired-end sequencing was performed using a NextSeq 500 sequencer (Illumina) with read lengths of 75 nt resulting in mean fragment lengths of around 200 nucleotides.

### Statistics

Statistically significant differences were determined by the two-tailed Student’s *t*-test where indicated. For natural transformation assays, data were log-transformed (62) before statistical testing. When the number of transformants was below the detection limit, the value was set to the detection limit to allow for statistical analysis.

### Other methods

Detailed information on strain design through recombinant DNA techniques and bioinformatic analyses are available in the *SI Appendix* under *Material and Methods*.

### Data availability

WGS reads of the 360 transformants have been deposited in NCBI’s Sequence Read Archive (SRA) under SRA accession numbers SRR6934824 to SRR6935183 according to *SI Appendix*, Table S3. The Bioproject accession number is PRJNA447902.

## Acknowledgement

The authors thank members of the Blokesch laboratory and F. Le Roux for discussions and A. Boehm for strain Sa5Y. We also acknowledge preliminary bioinformatic analyses by S. Strempel (Microsynth), A.-C. Portmann, and I. Mateus, who also uploaded the sequencing reads to NCBI. This work was supported by EPFL intramural funding, the Swiss National Science Foundation grant 31003A_162551, and a Starting (309064-VIR4ENV) and Consolidator (724630-CholeraIndex) grant from the European Research Council to MB. M.B. is a Howard Hughes Medical Institute (HHMI) International Research Scholar (grant #55008726).

## Author contributions

N.M. and M.B. designed research; N.M., S.S, C.S, and M.B. performed wetlab experiments; N.G. and C.I. performed bioinformatic analyses; N.M., N.G., C.I., and M.B. discussed the bioinformatic data; M.B. wrote the manuscript with input from N.M., N.G., and C.I. All authors approved the final version.

The authors declare no conflict of interest.

## Abbreviations

HGT –: horizontal gene transfer;
gDNA –: genomic DNA;
tDNA –: transforming DNA;
QS –: quorum-sensing

