## Supplemental information for "Neighbor predation linked to natural competence fosters the transfer of large genomic regions in *Vibrio cholerae*"

***SI - text***

**Long-read sequencing of O1 El Tor strain N16961 revealed additional rRNA clusters**

As a recent study identified a large inversion close to the origin of replication of chr 1 in the original genome sequence of N16961 (1), which most likely resulted from an imperfect assembly or a lab domestication event, we sequenced and *de novo* assembled the genome of our laboratory stock of strain N16961 (2). This stock is resistant to streptomycin, which is consistent with most literature reports on strain N16961, while the original reference strain N16961 remained sensitive to streptomycin according to its genome sequence. This difference suggests that mutagenesis event(s), including the characteristic streptomycin-resistance mutation within *rpsL* (encoding for RpsL[K43R]) must have occurred while the strain was domesticated. Comparative genomics between our laboratory stock of N16961 and the reference genome (3) showed a high level of sequence identity, though also confirmed the previously reported inversion around the origin of chromosome 1 (1) (Fig. S1). Additionally, significant differences were observed between the two N16961 genome sequences with respect to the number and arrangement of the ribosomal RNA clusters, which could have resulted from assembly artifacts. Indeed, while Heidelberg and colleagues described the presence of eight rRNA operons (16S-23S-5S) (3), we found ten rRNA operons (plus an additional 5S copy close to the tRNA-Thr). The only other major differences observed between the strains were in genes VC1620 and *vasX* (VCA0020) on chromosome 1 and 2, respectively, as mentioned in the main text (Fig. S1; marked with * and #). VC1620, or *frhA,* on chr 1 encodes a large protein with several cadherin tandem repeat domains, the number of which vary among different *V. cholerae* strains (4). Due to the repetitive nature of this DNA region, an assembly mistake in either of the two genome assemblies can therefore not be excluded. The sequence of the newly sequenced stock of N16961 was identical to the region in pandemic strain A1552 (Fig. S2). For the discrepancy within *vasX* on chr 2, we observed a significant number of single nucleotide polymorphisms (SNPs; n = 11) or nucleotide insertions (n = 14) within this 3,272-bp-long gene. One of these nucleotide insertions caused a frameshift that therefore resulted in a premature stop codon. As BLAST analyses suggested that this change was strain specific and not previously observed, we Sanger sequenced the corresponding DNA region using the same genomic DNA preparation that was used for the long-read PacBio sequencing, which allowed us to elucidate whether the mutations reflected a lab domestication event or whether the PacBio sequence was of low quality in this specific area of the genome. The latter turned out to be the case, as the mutations described above were absent from the Sanger sequencing reads. We therefore conclude that the two genomes are highly identical and that the major differences involve imperfect rRNA cluster assembly in the reference genome, which resulted in the underestimation of the number of rRNA clusters and the inverted assembly around the origin of replication of chr 1.

**Comparative genomics between pandemic strains N16961 and A1552**

*V. cholerae* O1 El Tor (Inaba) strain A1552 (formerly known as 92A1552-Rif$^r$) is a rifampicin-resistant derivative of strain 92A1552 (5), which was isolated in California from a traveller returning from South America. Epidemiological investigations concluded that the transmission of this strain occurred via a contaminated seafood salad that was served on an airplane between Lima, Peru and Los Angeles, California (6, 7), which links this strain to the Peruvian cholera outbreak in the 1990s.

Comparing the *de novo* assembled genome sequence of A1552 (2) to the genome of our laboratory stock of N16961 showed a large degree of genomic conservation between both isolates despite the fact that strain A1552 contained only nine rRNA operons (16S-23S-5S) instead of the ten operons in strain N16961. The genome architecture, however, was not maintained, as strain A1552 had undergone a large inversion of approximately 2.7 Mbp between two rRNA clusters on chr 1 (Fig. S2; genome sequence inverted to simplify visualization), a finding that is consistent with a previous report (8). Apart from this inversion, the major differences between both strains were the presence of the WASA-1 island (a 44-gene-carrying element of unknown function; (9)) and a modified *Vibrio* seventh pandemic island II (VSP-II) in strain A1552 (Fig. S2). These features allowed us to classify strain A1552 as belonging to the LAT-1 sublineage of the West-African South American (WASA) lineage of the seventh pandemic *V. cholerae* strains (9, 10).

### *SI - Materials and Methods*

**Bacterial strains and growth conditions.** Bacterial strains and plasmids used in this study are described in Table S1. Bacteria were routinely grown aerobically in lysogeny broth (LB) or on LB agar plates (1.5% agar) at 30°C or 37°C. Half-concentrated defined artificial seawater medium (0.5x DASW) containing HEPES and vitamins (11) was used for chitin-induced T6SS killing and natural transformation experiments. Agar plates containing M9 minimal medium (Sigma-Aldrich) supplemented with vitamins (MEM vitamin solution; Gibco), 0.001% casamino acids (Merck), and 0.2% mannose were used to select *V. cholerae* strain A1552 and to exclude strain Sa5Y (to check the direction of transformation for *comEC*-positive prey). Thiosulfate-citrate-bile salts-sucrose (TCBS; Sigma-Aldrich) agar plates were used to counterselect *E. coli* strains after mating with *V. cholerae*. Antibiotics were used at the following concentrations whenever required: chloramphenicol (Cm), 2.5 μg/ml; kanamycin (Kan), 75 μg/ml; streptomycin (Strep), 100 μg/ml; ampicillin (Amp), 100 μg/ml; and rifampicin (Rif), 100 μg/ml.

**DNA manipulation techniques.** Recombinant DNA techniques were performed following standard molecular–biology-based protocols (12). DNA-modifying enzymes such as Pwo DNA polymerase (Roche), Taq DNA polymerase (GoTaq; Promega), and restriction modification enzymes (New England Biolabs) were used according to the manufacturer's recommendations. Genetically modified strains were verified by colony PCR and, if required, also by Sanger sequencing (Microsynth, Switzerland) for their correctness.

**Genetic engineering of bacterial strains.** To delete gene(s) from the parental WT strains (A1552 or Sa5Y), a gene-disruption method based on either a counter-selectable suicide plasmid pGP704-Sac28 (13) or on natural transformation and FLP recombination was used (TransFLP method; (14-16)). Natural transformation was also used to insert the antibiotic resistance cassettes *aph* ($Kan^R$), *cat* ($Cm^R$), and/or *bla* ($Amp^R$) into target gene(s) of *V. cholerae*.

**Genome comparisons.** Each chromosome was segmented in contiguous fragments of 10 kb, which were locally aligned against the corresponding chromosome of a reference genome. Each fragment was aligned in the forward and reverse orientation, and the best alignment was retained.

The number of differences per 10 kb was evaluated by counting the number of events necessary to mutate the reference to obtain the 10 kb fragment (e.g. an insertion or deletion of an arbitrary number of nucleotides would count as one event). To visualize the overall architecture and differences, circular plots in which each 10 kb fragment was linked to its reference genome counterpart and colored according to the number of differences were made in R using the circlize package (17). Black and blue linkers indicated whether the 10 kb fragment had the same or reverse orientation relative to the reference.

**Scoring of horizontal gene transfer events through bioinformatics analyses.** HGT events were scored for the 360 transformants that were derived from the eight different experimental conditions (Fig. 4 and listed in Table S2). For each condition, a predator/acceptor strain (A1552 or its derivative) and a prey/donor strain (derivatives of Sa5Y; used in mixed cultures or as purified gDNA) were defined, and their genomic sequences were generated *in silico*. These *in silico* templates were based on the recently announced genome sequences of the parental strains (2) to which the integrated genomic features were added (e.g., integration of *aph*, *cat*, and/or *bla* cassettes proceeded by constitutive promoters). Each DNA template contained two parts reflecting the large ~3.0 Mbp chr 1 and the small ~1.1 Mbp chr 2. Preliminary analyses identified the presence of systematic differences in each sample, which can be attributed to errors in the reference templates. Thus, before starting the final analysis process, the following patches were applied to the chr 2 of the predator/acceptor reference genomes:

- Coordinate (in reference genome CP028895): 445515–445520 => TTTTTT replaced by TTTTT.
- Coordinate (in reference genome CP028895): 447288–447289 => GC replaced by G.
- Coordinate (in reference genome CP028895): 467939–467940 => AT replaced by A

(resulting in a replacement of TTTTTTT [467940–467946] by TTTTTT). The correctness of these *in silico* changes was confirmed by Sanger sequencing.

The FASTA sequences of the corrected reference templates used for this work are available in the Reference directory on GitHub: https://github.com/sib-swiss/VibrioCholerae_HGT.

Each read pair was aligned against both genomes of the donor and acceptor strains, and the position with the least number of mismatches was kept. If multiple possibilities with the same number of mismatches were possible, the possibilities were kept for later processing. The alignments were performed with an in-house code derived from the fetchGWI tool [https://sourceforge.net/projects/tagger/files/fetchGWI-tagger/], but other tools such as BWA or bowtie2 would equally qualify for the same purpose. The results of this first analysis phase were obtained as tab-delimited lists of read pairs with their alignment positions, number of mismatches for each of the reads of the pair, and number of matching positions with the same number of mismatches.

The second step of the analysis consisted of a C program that parsed the tab-delimited file of the previous step and recorded the accumulated coverage and the observed nucleotide for each position of each of the donors' and acceptors' chromosomes. For each analysis, two separate recordings were kept: the first recording took only those read pairs into account that showed zero mismatches together with a unique unambiguous matching position, while the second recording took all other matches into account (e.g., a coverage of 1 was counted in each matched position regardless of whether the match was unique or not). At the end of this process, regions with continuous coverage were created by looking for positions in the first recording (unique and exact matches) for which the observed coverage was at least 3. The start and end positions of such continuous stretches were determined by extending as far as possible from this seed position, taking into account the total coverage from both recordings. If the length of the so-defined fragment was at least 500 nucleotides it was kept for output. The output consisted of a

FASTA file containing all the defined fragments and the FASTA header of each fragment. This header recorded which reference was covered (e.g., which strain [acceptor or donor] and which chromosome [large chr 1 or small chr 2]). For each position, the nucleotide that composed more than half of the coverage of that position was generated in the output. If no single nucleotide represented more than half of the coverage, an N was used.

The expectation was that the fragments obtained in the previous step constituted a full coverage of the transformant, which had inherited its genomic material mostly from the parental acceptor strain with some DNA regions originating from the donor strain by HGT; this referred particularly to those regions containing the antibiotic resistance cassette, which was used as selective marker. A global DNA alignment for each output fragment was therefore performed by mapping the fragments onto the corresponding chromosomes of the donor and acceptor strains. Since we determined that the donor and acceptor genomes differed on average by 1 nucleotide every 55 nucleotides, it was expected that several mismatches would be found when a fragment originating from the donor strain was aligned to the genome of the acceptor strain. The first and last mismatches of those fragments were defined as outer bounds of the transferred fragments, as they arguably represented the minimal length of the transferred DNA region. The summary data of these alignments and boundary information were determined using Perl scripts as were those regions of the acceptor genomes that were not covered by reads derived from the analyzed transformant (e.g., transformed acceptor strain). The output of those scripts (available on GitHub: https://github.com/sib-swiss/VibrioCholerae_HGT) was then used to generate a final table of transferred segments as well as summary plots by applying the R package circlize (17). Finally, the data were transferred to Excel to calculate the total length of horizontally acquired DNA per transformant as well as the number of HGT events per transformant. The GraphPad Prism software was used for graphic visualization. To validate the bioinformatic approach, the mapped reads obtained for >50 samples were also visually inspected for transferred regions using the software Geneious®.

***SI – References***

12. Sambrook J, Fritsch EF, & Maniatis T (1982) *Molecular Cloning: A Laboratory Manual*, (Cold Spring Harbor Laboratory Press, Cold Spring Harbor, NY.).

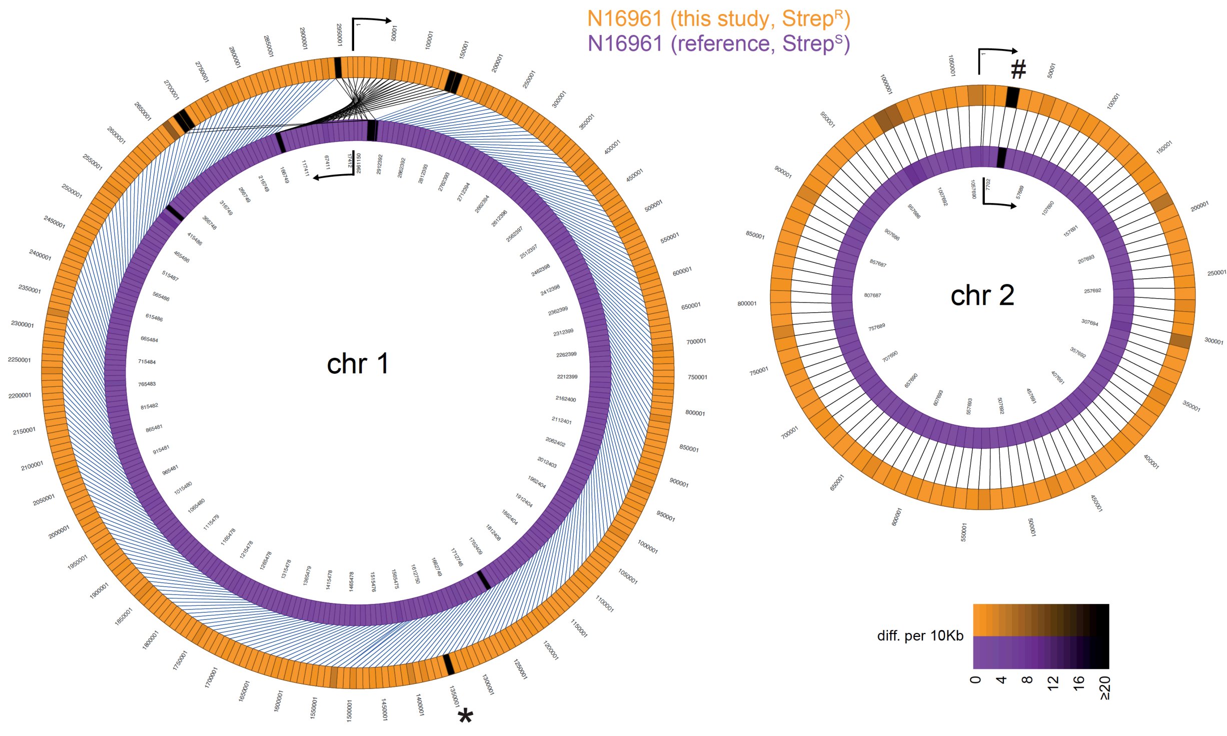


**Figure S1: Comparative genomics of *V. cholerae* reference strain N16961 and a newly sequenced laboratory stock of the same strain.** The newly sequenced genome of the streptomycin-resistant laboratory stock of strain N16961 (long-read PacBio sequencing technology; (2)) was compared to the genome sequence of the reference genome (Heidelberg *et al.*, 2000). Detailed explanations on the comparison are as described for Figure 2. Marked discrepancies are based on repetitive sequences (*) or a sequencing error (#) as discussed in the *SI Appendix*.

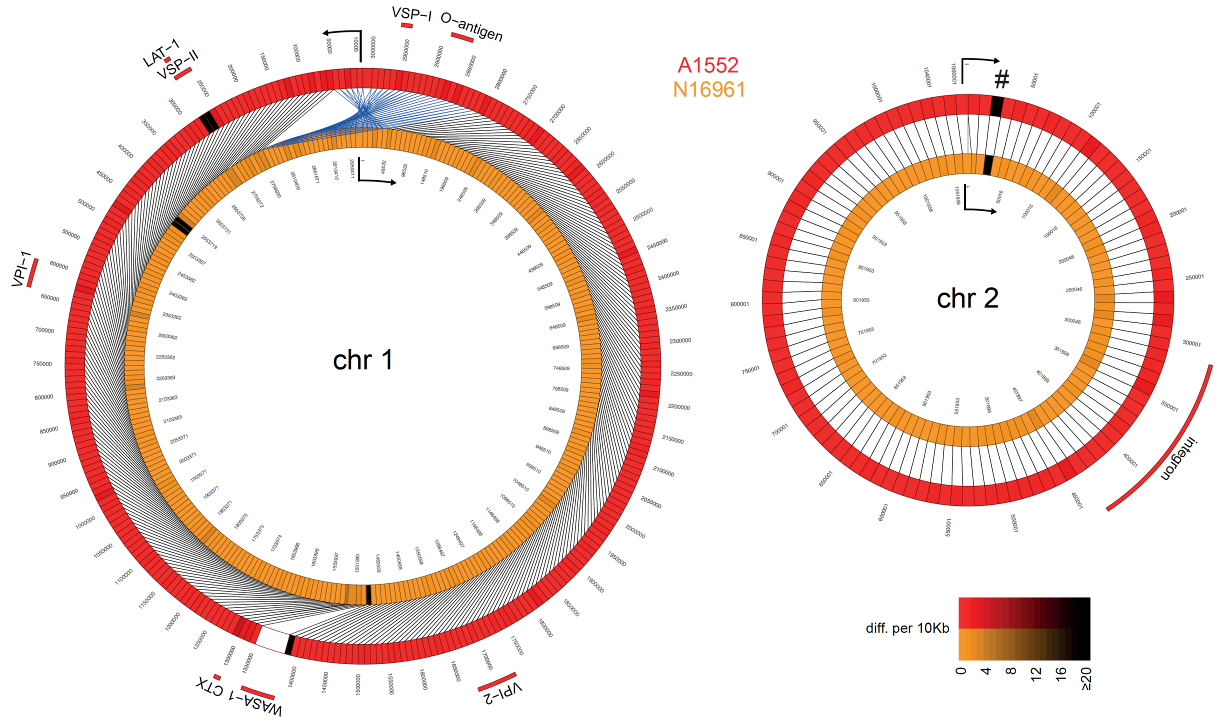


**Figure S2: Comparative genomics of pandemic *V. cholerae* strains N16961 and A1552.** The genome sequences of our laboratory stock of pandemic strains N16961 and A1552 were compared. Details are as described in Figure 2. The marked discrepancy (#) resulted from a sequencing error, as discussed in the *SI Appendix*.

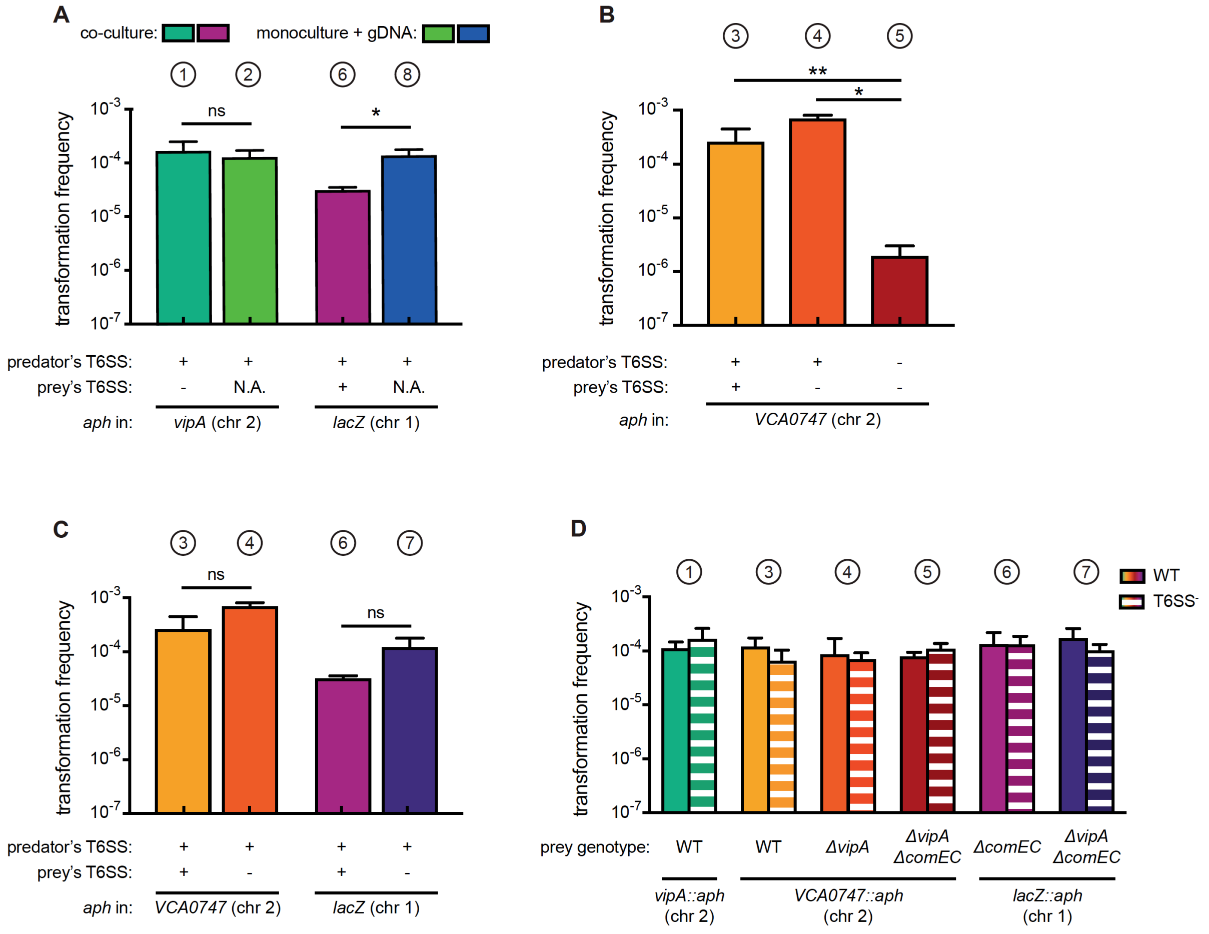


**Figure S3: Natural transformation is enhanced by T6SS-mediated killing of prey bacteria.** (*A-C*) Transformation is enhanced in T6SS-positive predator cells. To induce natural competence, cultures were grown on chitin flakes. Bacteria were grown as co-cultures (predator + prey) or as monocultures (predator only). In the latter case, purified gDNA served as the transforming material. Transformants were selected based on their acquisition of the *aph* resistance cassette located in *vipA*, *VCA0747,* or *lacZ* on chr1 or chr2, as indicated below the graphs. The killing capability of each strain is indicated below the graph (e.g., T6SS + or -). Transformation frequencies are shown on the Y-axis (± SD, as depicted by the error bars) and depict averages of at least two biologically independent experiments (corresponding to the experiments described in Figure 4 and maintaining the same color code). N.A., not applicable. (*D*) The location of the resistance genes does not influence the transformation efficiency. Natural transformability of the WT (plain bars) or its T6SS-minus derivative (T6SS$^-$; Δ*vipA;* hashed bars) was scored using gDNA as the transforming material. The gDNA samples were derived from the prey strains of conditions ① and ③–⑦ and the respective genotype is shown below the graph. Transformation frequencies were scored based on the acquisition of the *aph* resistance cassette, which was integrated into different genes on chromosome 1 or 2 (chr 1/chr 2). Data represent the average of three independent experiments (±SD). (*A-D*) Statistical significance is indicated (*$p < 0.05$; **$p < 0.01$).

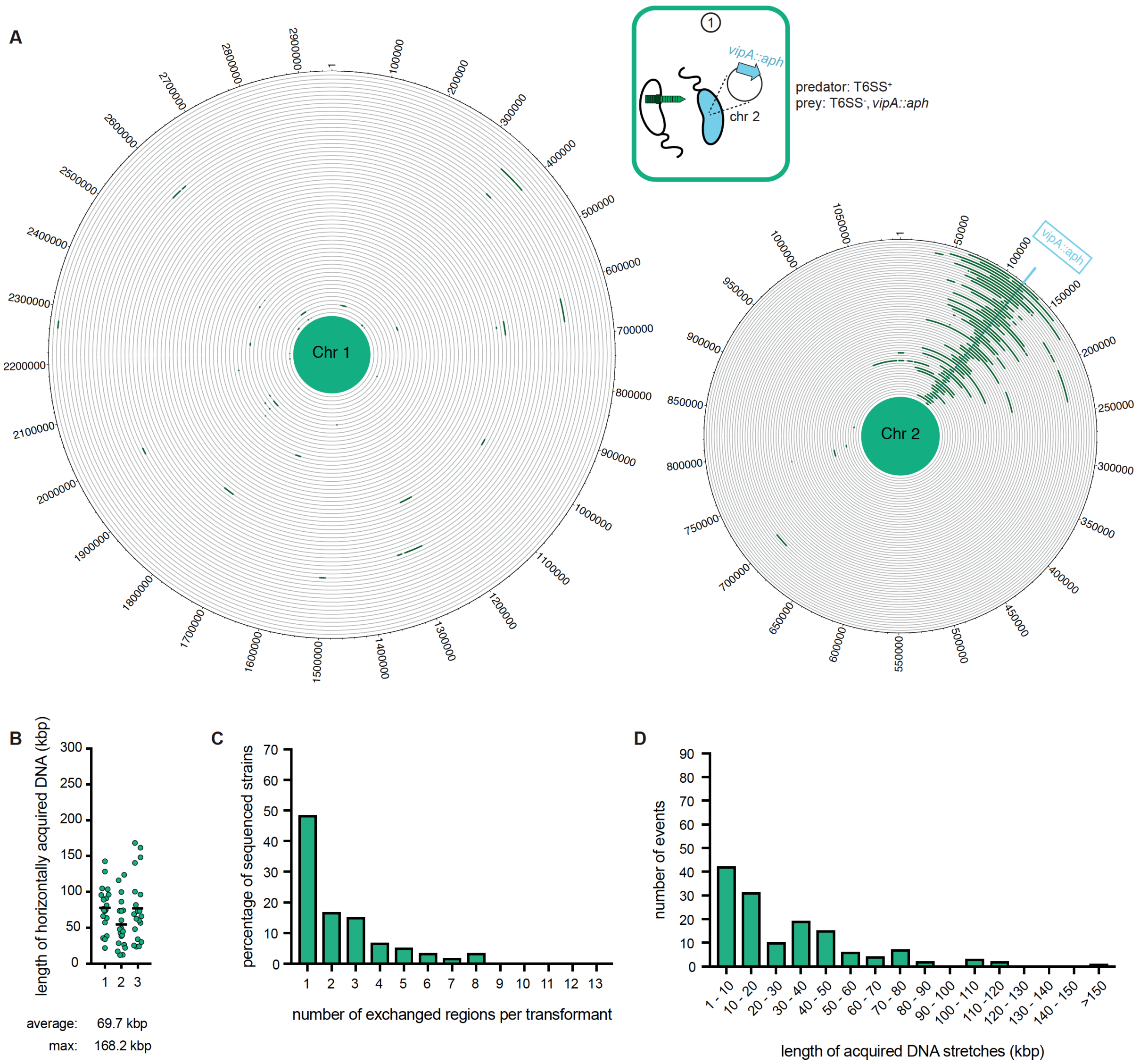


**Figure S4: WGS-based quantification of horizontally acquired DNA under condition ①.** Data as in Figure 3 with the addition of the map of both chromosomes in panel A.

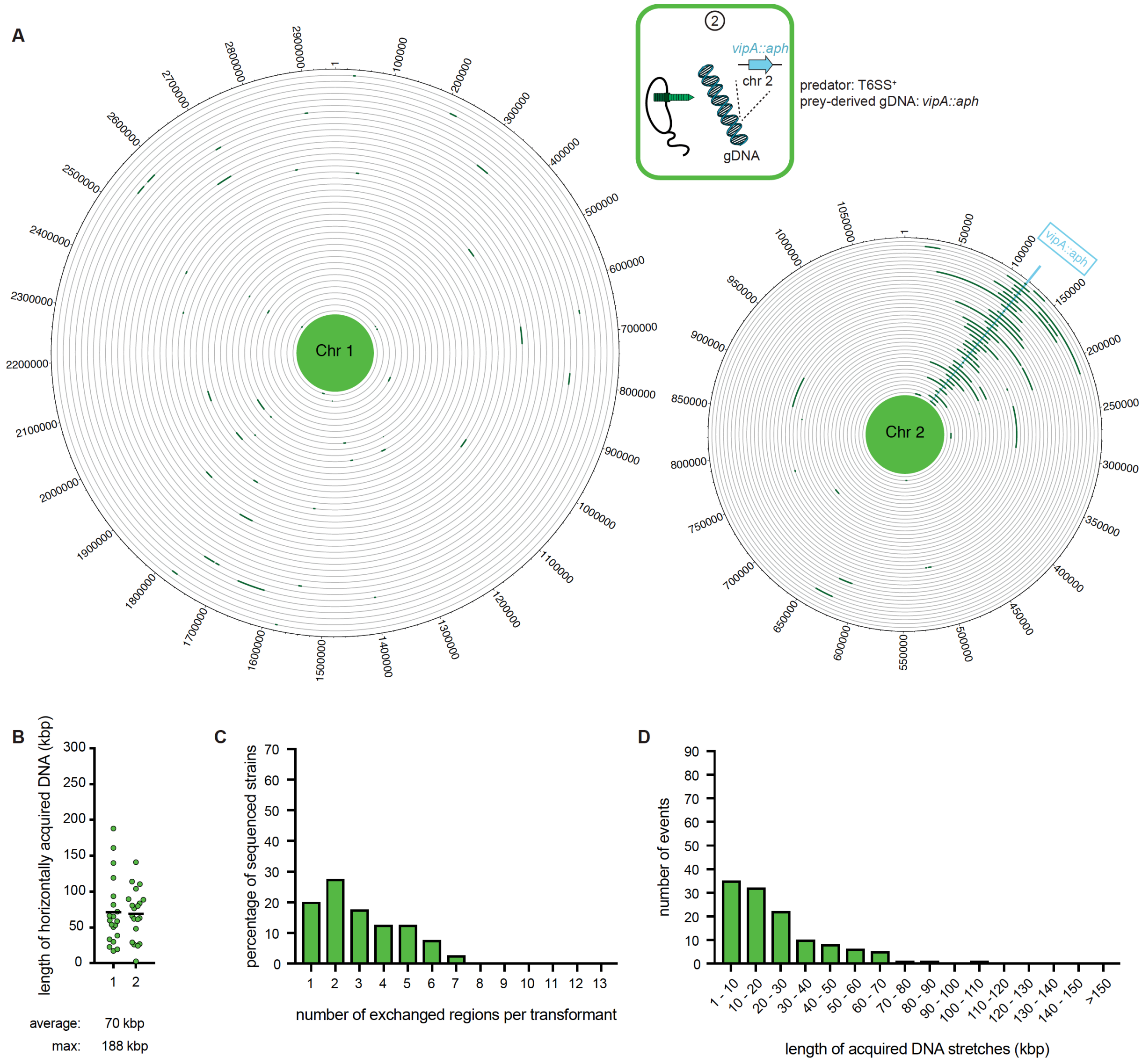


**Figure S5: WGS-based quantification of horizontally acquired DNA under condition ②.** Details as described for Figure 3 with the addition of the map of both chromosomes in panel A.

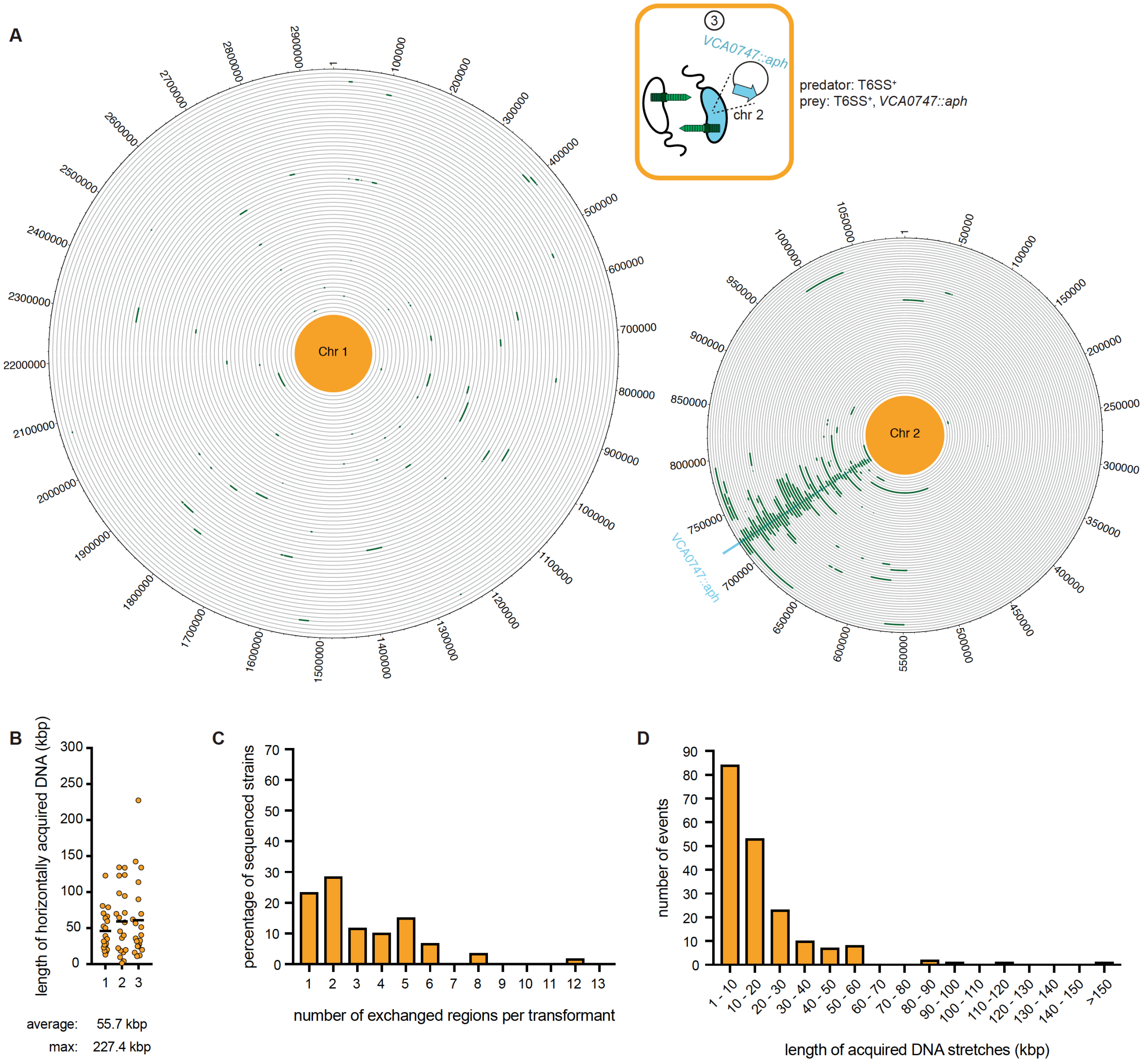


**Figure S6: WGS-based quantification of horizontally acquired DNA under condition ③.** Details as described for Figure 3 with the addition of the map of both chromosomes in panel A.

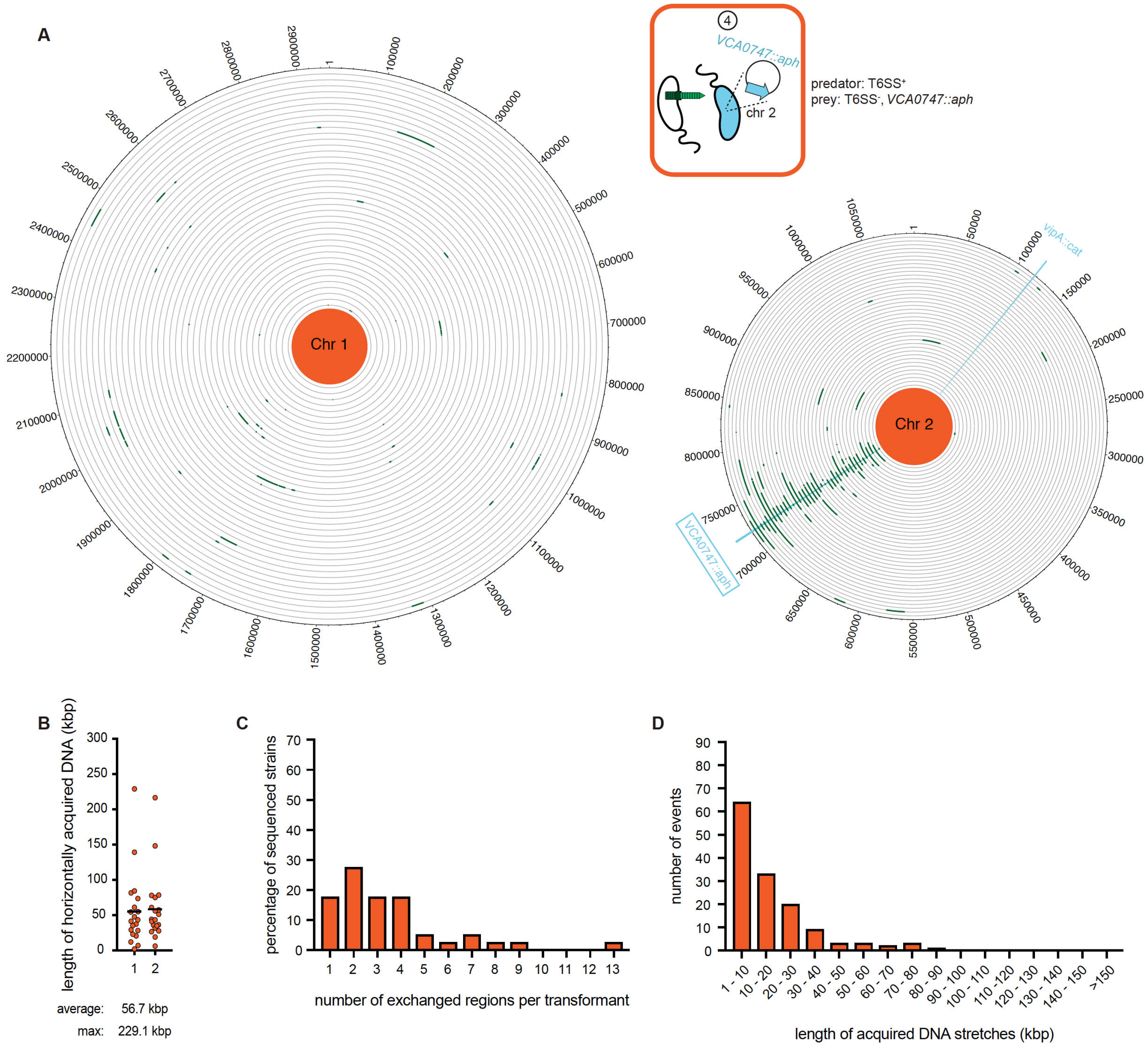


**Figure S7: WGS-based quantification of horizontally acquired DNA under condition ④.** Details as described for Figure 3 with the addition of the map of both chromosomes in panel A.

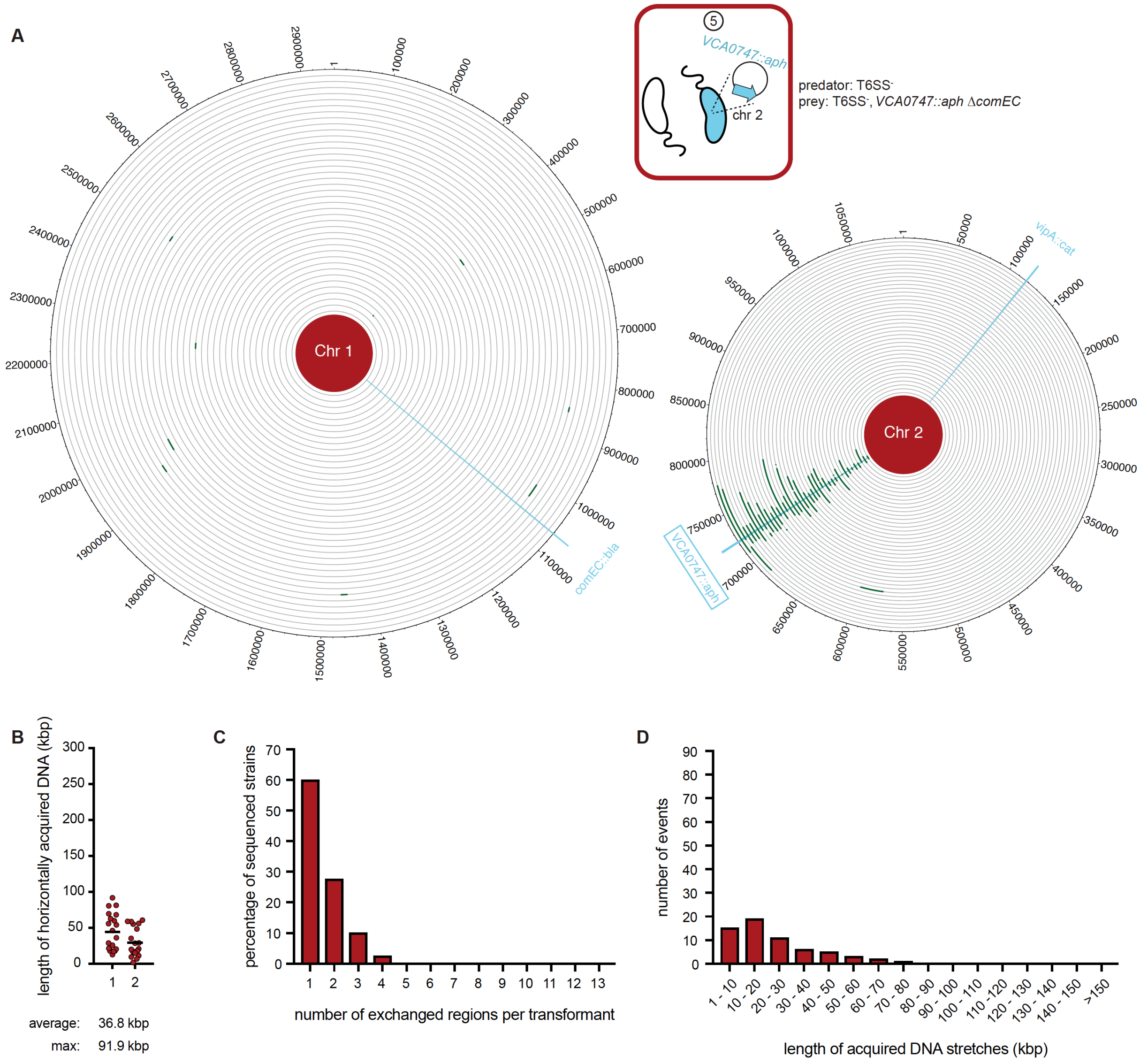


**Figure S8: WGS-based quantification of horizontally acquired DNA under condition ⑤.** Details as described for Figure 3 with the addition of the map of both chromosomes in panel A.

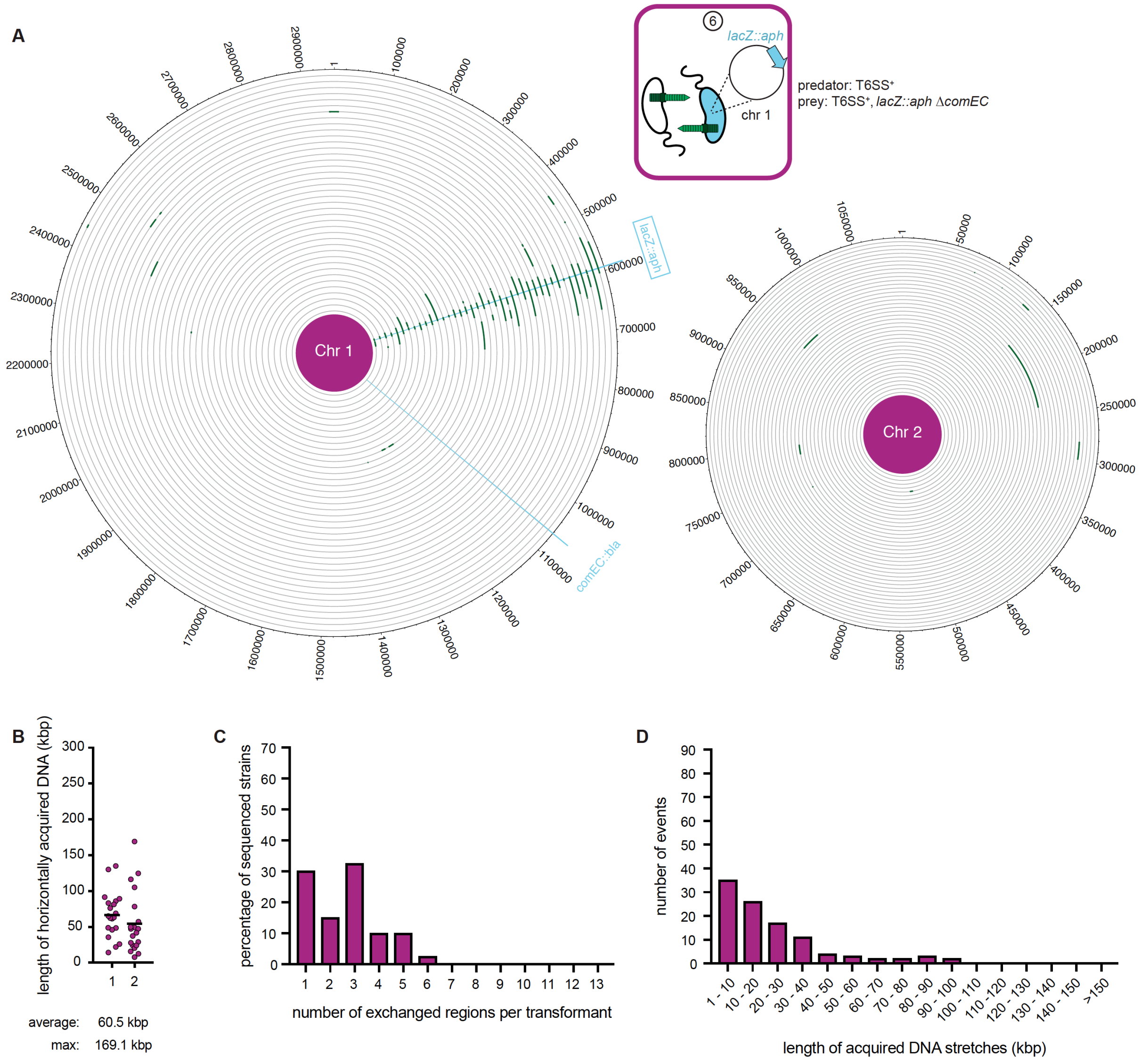


**Figure S9: WGS-based quantification of horizontally acquired DNA under condition ⑥.** Details as described for Figure 3 with the addition of the map of both chromosomes in panel A.

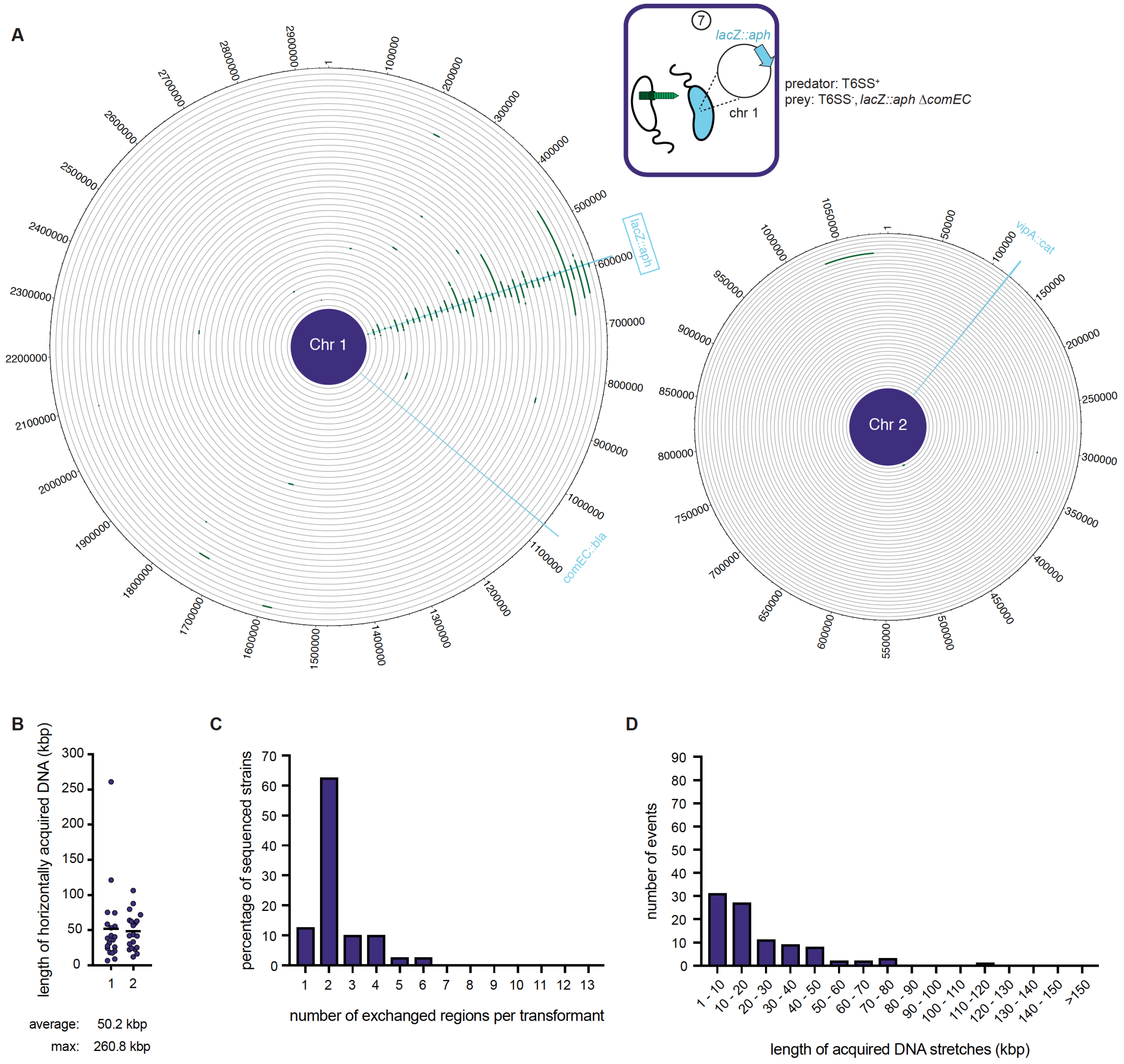


**Figure S10: WGS-based quantification of horizontally acquired DNA under condition ⑦.** Details as described for Figure 3 with the addition of the map of both chromosomes in panel A.

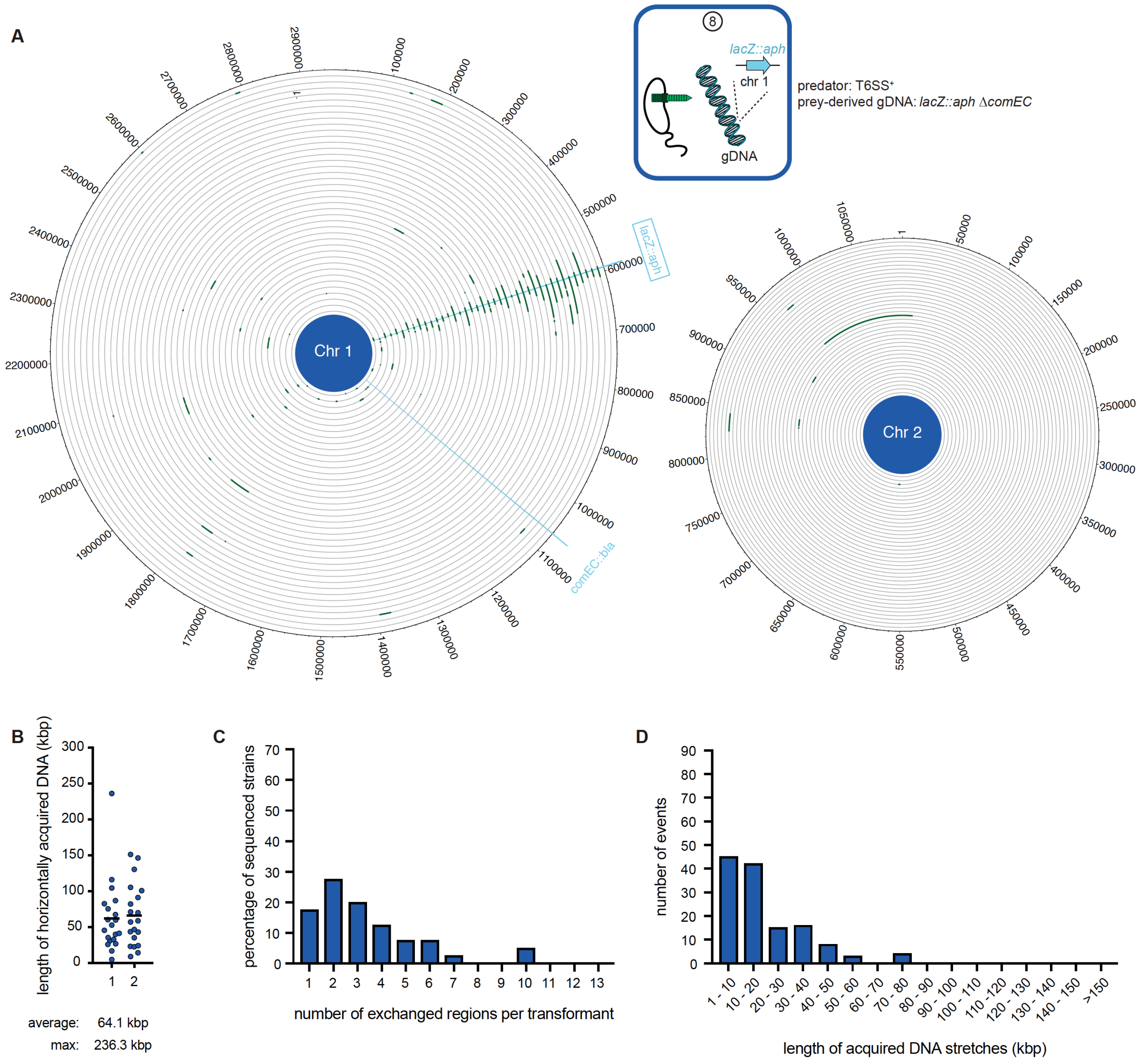


**Figure S11: WGS-based quantification of horizontally acquired DNA under condition ⑧.** Details as described for Figure 3 with the addition of the map of both chromosomes in panel A.
